# Precise hyperacuity estimation of spike timing from calcium imaging

**DOI:** 10.1101/790600

**Authors:** Huu Hoang, Masa-aki Sato, Shigeru Shinomoto, Shinichiro Tsutsumi, Miki Hashizume, Tomoe Ishikawa, Masanobu Kano, Yuji Ikegaya, Kazuo Kitamura, Mitsuo Kawato, Keisuke Toyama

## Abstract

Two-photon imaging is a major recording technique in neuroscience, but it suffers from several limitations, including a low sampling rate, the nonlinearity of calcium responses, the slow dynamics of calcium dyes and a low signal-to-noise ratio, all of which impose a severe limitation on the application of two-photon imaging in elucidating neuronal dynamics with high temporal resolution. Here, we developed a hyperacuity algorithm (HA_time) based on an approach combining a generative model and machine learning to improve spike detection and the precision of spike time inference. First, Bayesian inference estimates the calcium spike model by assuming the constancy of the spike shape and size. A support vector machine employs this information and detects spikes with higher temporal precision than the sampling rate. Compared with conventional thresholding, HA_time improved the precision of spike time estimation up to 20-fold for simulated calcium data. Furthermore, the benchmark analysis of experimental data from different brain regions and simulation of a broader range of experimental conditions showed that our algorithm was among the best in a class of hyperacuity algorithms. We encourage experimenters to use the proposed algorithm to precisely estimate hyperacuity spike times from two-photon imaging.

## Introduction

Recently, two-photon imaging has been one of the major means of recording multineuronal activities in neuroscience to obtain the precise morphology and location of the target neurons because of its high spatial resolution^1–6^. However, its utility is still constrained by its relatively low temporal resolution due to the mechanical scanning of two-photon rays. The other problems are the nonlinearity, the slow dynamics and the low signal-to-noise ratio (SNR) of the calcium (Ca) responses^7–10^. Many algorithms have been proposed to reconstruct spike trains from Ca imaging data, including conventional thresholding^11^, deconvolution^12–15^, template matching^16–20^, Bayes inference^21–23^ and machine learning^24,25^, to overcome these problems. Few of them, however, have addressed the two challenging goals simultaneously: reliable spike detection and spike time estimation with high temporal precision in the presence of the nonlinearity, slow dynamics and low SNR of the Ca responses^26^. For the former goal, the spike dynamics of the target neurons and/or kinematics of the Ca responses may vary dramatically across brain regions and different Ca dyes. For the latter goal, a trade-off between the number of recorded neurons and temporal resolution exists. The slow kinematics and the low SNR of the currently available Ca dyes may also limit the temporal precision of the information conveyed by the Ca responses. These factors impair reliable spike detection as well as precise spike time estimation for high-frequency firing that is frequently encountered in cortical cells^27–29^.

Here, we propose an approach combining a generative model of Ca responses including nonlinearity and dye dynamics with a supervised classifier to overcome the aforementioned difficulties. Our hyperacuity algorithm, named HA_time (HyperAcuity time estimation), estimates the Ca spike model by Bayesian inference assuming size and shape constancy, compensates for the nonlinearity of the Ca responses, and detects spikes from Ca imaging data by a support vector machine (SVM) using the ground-truths, i.e., simultaneously recorded electrical spikes, as supervised information. To achieve hyperacuity precision, spike timings were calibrated to minimize the residual errors in model prediction using the hyperacuity vernier scale. This approach benefits from the advantages of both generative models and supervised learning. On the one hand, the Ca spike model is utilized to provide supplemental information for spike detection as well as to estimate the spike times at higher temporal precision than the sampling resolution. On the other hand, the supervised learning compensates for fluctuations in the Ca responses due to noise and sampling jitters, which are not considered by the generative model. As a consequence, HA_time can improve both the spike detection and spike time estimation of two-photon recordings.

A simulation study confirmed that compared with the thresholding algorithm, HA_time improved the temporal precision by 2-20-fold. The previous algorithms have aimed to improve spike detection as well as spike time estimation with higher temporal precision than that expected for the sampling rate of two-photon recordings. They assumed generative models for spike generation and maximized the likelihood of the estimates^17,21,22^. Accordingly, hyperacuity performance was limited to only the cases where the Ca responses satisfy the assumptions of generative models. To prove the advantages of the approach combining the generative model and supervised learning, we compared our method with four previous hyperacuity algorithms^16,17,22,23^. The benchmark results for the experimental data sets showed that HA_time was among the best across three brain regions: the cerebellar, hippocampal and visual cortices. Furthermore, the simulation analysis conducted across a broad range of parameters for the experimental conditions, including the mean neuronal firing frequency, the nonlinearity, the decay time of the Ca dyes, and the sampling rate, providing useful information for users to select the most suitable algorithms for the given experimental conditions and highlighting the advantages of our algorithm over the other ones under the high firing frequency and/or strong nonlinearity conditions frequently encountered in the cortical cells of behaving animals.

## Results

### Hyperacuity algorithm for spike timing estimation

Our hyperacuity algorithm, HA_time, was conducted in three steps: 1) Bayesian inference of the Ca spike model from Ca imaging data, 2) spike detection by SVM assisted by the matching information of the Ca imaging data with the Ca spike model and 3) hyperacuity spike time estimation to minimize the errors between the Ca response model prediction and the recorded Ca imaging data using the hyperacuity vernier scales. We noted here that the term “Ca spike model” indicates the constant Ca transient (i.e., the amplitude and shape) of a single spike, whereas “Ca response model” is a generative model of the Ca spike model, superposition of multiple spikes, nonlinearity of the Ca responses and noise (see below).

We assumed that the Ca imaging data were sampled from the Ca spike model (double exponentials) with variable sampling jitters between the onsets of the Ca spike and the sampling times. The Ca spike responses were first linearly superimposed for multiple spikes in short intervals and added with the Gaussian noise. The sub- or superlinearity of the Ca responses was determined by comparing the observed Ca imaging data with the data predicted by the Ca response model. We compensated for the nonlinearity by inversely transforming the observed Ca imaging data by nonlinearity models fitted by logarithmic functions (Fig. 1A).

**Figure 1:**
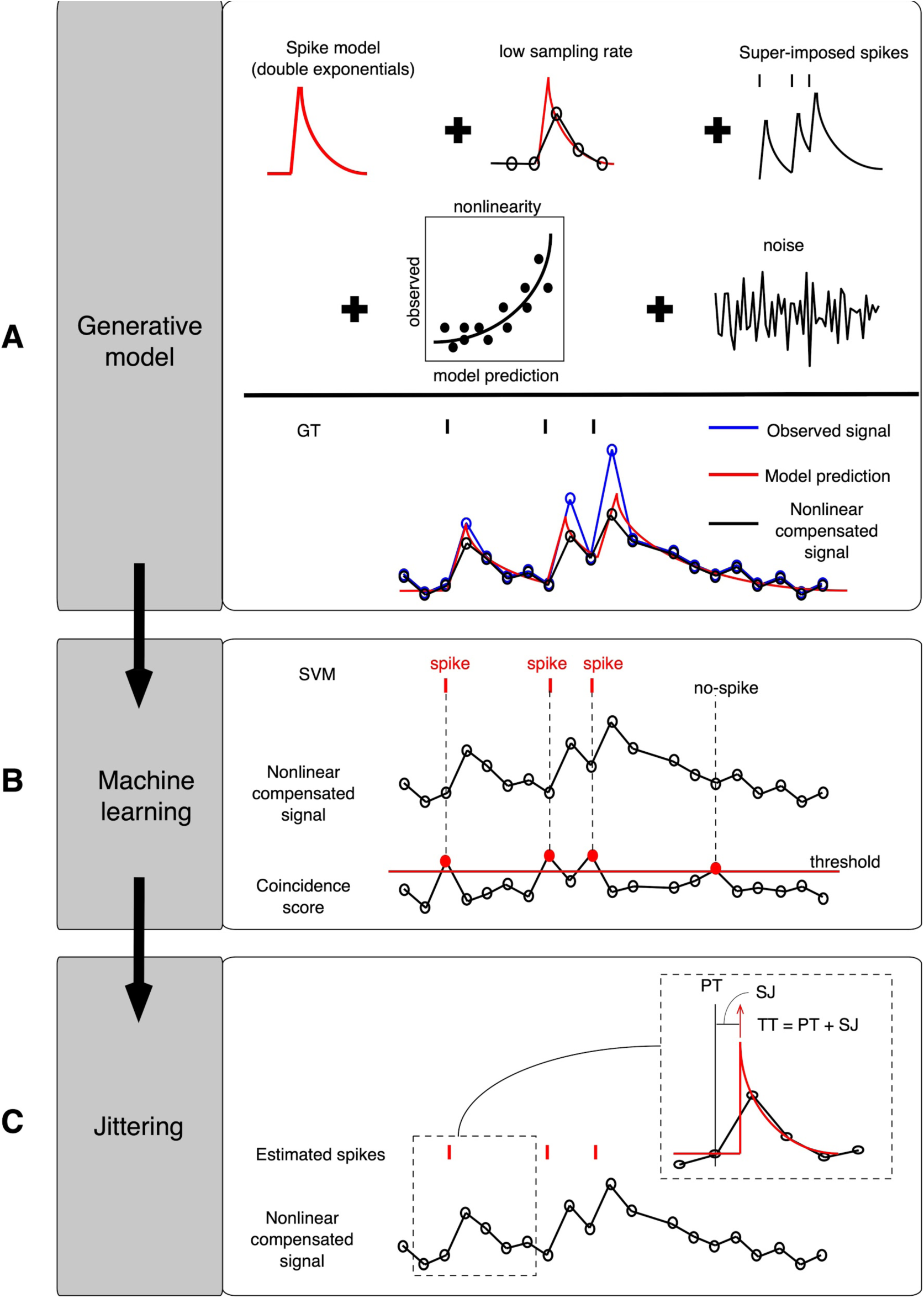
The hyperacuity support vector machine (HA_time) algorithm. A: The generative model assumed that Ca responses were sampled from the Ca spike model (double exponentials) with variable sampling jitters (SJs) due to the low sampling rate, superimposed by multiple spikes, fluctuated by nonlinearity and supplemented with Gaussian noise. Nonlinearity observed in the data was compensated by the nonlinearity model defined by logarithmic functions of the observed Ca imaging data and Ca response model prediction. B: The coincidence score, computed by convoluting the first derivative of Ca imaging data and that of the Ca spike model, was used to select spike candidates. Support vector machine (SVM) was trained to classify spikes and non-spikes from spike candidates with the Ca imaging data, the coincidence scores and the electrical spikes as feature, attribute, and teaching signals, respectively. C: The true spike time (TT) was estimated as the sum of SJ and pseudo-spike time (PT, the point that exceeds the threshold) minimizing the residual error of the Ca response model prediction using hyperacuity vernier 10-fold finer than the sampling interval.

Next, we estimated the coincidence score as a convolution of the first-order derivative of the Ca imaging data and that of the Ca spike model. A coincidence score threshold was used to sample the data segments as spike candidates. Here, the threshold and segment size were optimized to maximize the F1-score of the training data (see Methods). The SVM was trained to classify the sampled data segments into spike or non-spike segments. For this purpose, we fed the sampled Ca imaging data and the coincidence scores as the primary and attribute inputs, respectively, to the SVM and used the electrical spikes (ground truth) as the teaching signals (Fig. 1B).

For the test data, spike candidates were sampled in the same way as the training data, and the trained SVM detected the spikes among the candidates. We tentatively determined the time when the coincidence score exceeded the threshold as the pseudo-spike time (PT in Fig. 1C inset) and estimated the sampling jitters (SJs) to minimize the errors between the Ca response model prediction and the Ca imaging data using the hyperacuity vernier scales (10-fold finer time bin than the sampling interval). The true spike time (TT) was calculated as the sum of the PT and SJ (Fig. 1C).

### Hyperacuity improvement of HA_time in simulation

To illustrate the hyperacuity improvement of HA_time, we compared our method with the thresholding algorithm using the simulated data with no nonlinearity and fast Ca decay time (see Methods). Here, the threshold of the thresholding algorithm was optimized to maximize the F1-score of the training data.

Figure 2 shows the hyperacuity improvement, determined as the ratio of the mean of the spike time errors for the traditional thresholding algorithm to that for HA_time (see Methods), as a function of the mean firing frequency of the simulated spike train. Compared with the thresholding algorithm, in cases with a high sampling rate of 40-60 Hz, HA_time improved the temporal precision more than 5-fold under low firing rate conditions (<5 Hz) and maintained an approximately 2-fold improvement under high firing rate conditions, probably due to the decreased performance of both algorithms (thicker lines, Fig. 2). Such a tendency was also found in the cases with a lower sampling rate of 10-30 Hz, although the hyperacuity improvement was smaller (thinner lines). These results indicated that even under fairly simple data conditions, for which the conventional methods are widely used, HA_time was able to provide a 2-20-fold improvement in the temporal precision compared to that provided by the thresholding algorithm. For example, if we take an ideal case of a 60 Hz sampling frequency and 2 Hz firing frequency, the hyperacuity improvement was 12, meaning that the effective sampling rate is approximately 60 × 12 = 720 Hz, which is quite satisfactory.

**Figure 2:**
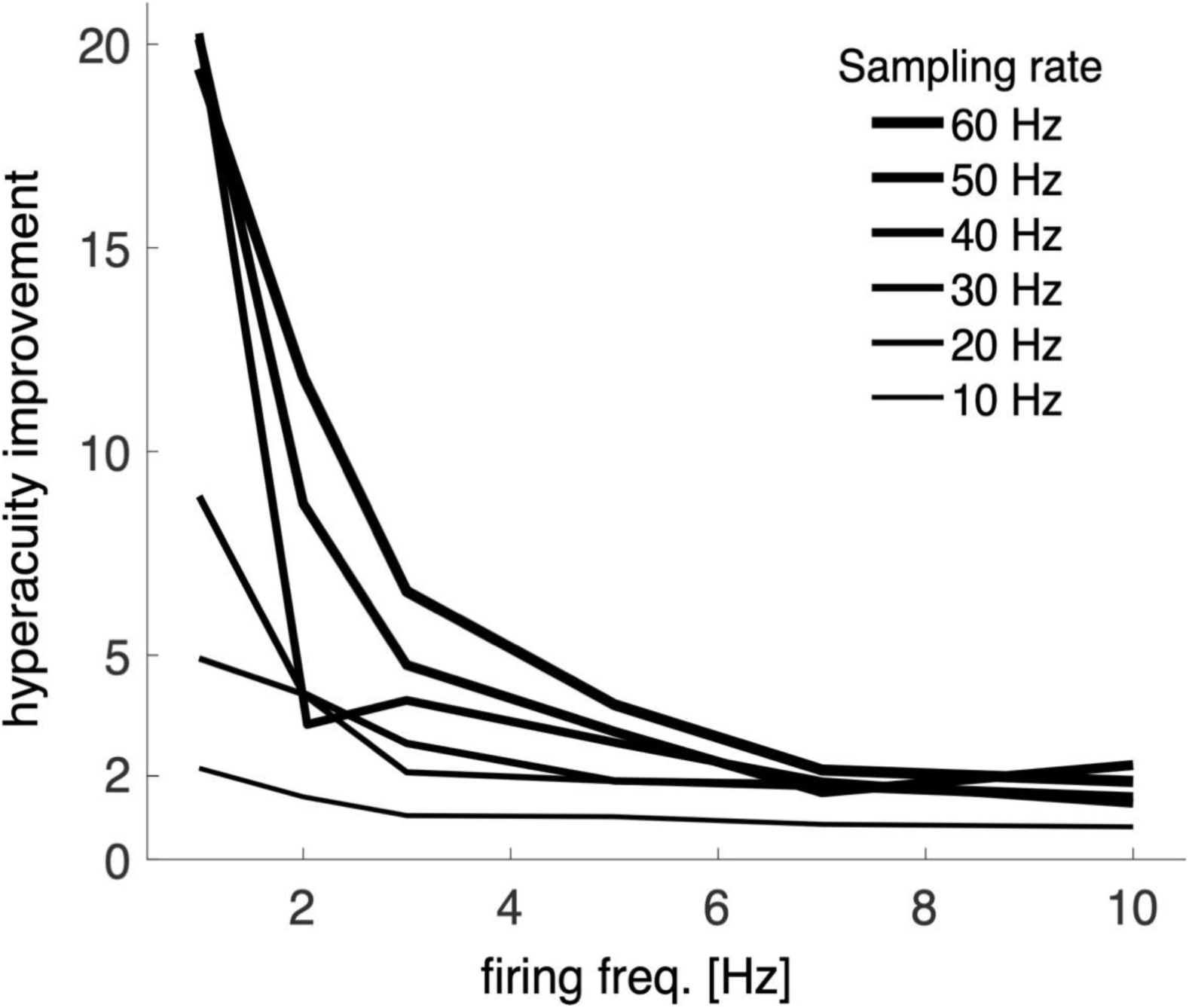
The hyperacuity improvement of HA_time. The hyperacuity improvement of HA_time compared with conventional thresholding as a function of the firing frequency at various sampling rates of 10-60 Hz (encoded by line thickness). The nonlinearity parameter α was fixed at 1, and τ_2_ and SNR were fixed at 0.2 s and 5, respectively.

### Application of HA_time to experimental data

We applied HA_time to noisy Ca imaging data obtained in cerebellar, hippocampal and primary visual cortical cells by two-photon recording with relatively low sampling rates.

For the two-photon recording of the Ca response from five Purkinje cells in the cerebellum (the dye, Cal-520, and sampling rate, 7.8 Hz), we sampled thirty-six data segments (segmental length, 2 s), each including a single electrical spike from the simultaneous electrical recording (sampling rate, 20 KHz), and constructed the Ca spike model by Bayesian inference. In agreement with the assumption of spike-shape constancy, the Ca spike model (τ_1_ = 0.05 s, τ_2_ = 0.4 s, red trace) was slightly faster in the rise time (red trace of Fig. 3A) than the electrical spike-triggered average of the Ca imaging data, while the amplitude of the Ca spike model roughly agreed with those for the spike-triggered averages. The longer time course of the spike-triggered response may be due to the sampling jitters. We also conducted a Bayesian estimation of the Ca spike models for the data from the entire hippocampus (n=9 cells, sensor, OGB1-AM and sampling rate, 10 Hz) and visual cortex (n=11 cells, sensor, GCAMP6f, sampling rate of 60 Hz). We avoided segments that contain burst activity (interspike interval < 2 s) since the Ca responses in the hippocampus and visual cortex showed strong nonlinearity during the burst activity (see below). A similar tendency to that in the cerebellum data was also noticed in the Ca spike model (red traces, Figs. 3C&E), which was faster in the rise time than the spike-triggered averages (blue traces) for both the hippocampus and visual cortex data. The dynamics of the dyes estimated by our Bayesian method were in agreement with those reported in previous studies (τ_1_ = 0.01 and τ_2_ = 0.2 s for GCAMP6f^30^ and τ_1_ = 0.1 and τ_2_ = 0.75 s for OGB1-AM^11^).

**Figure 3:**
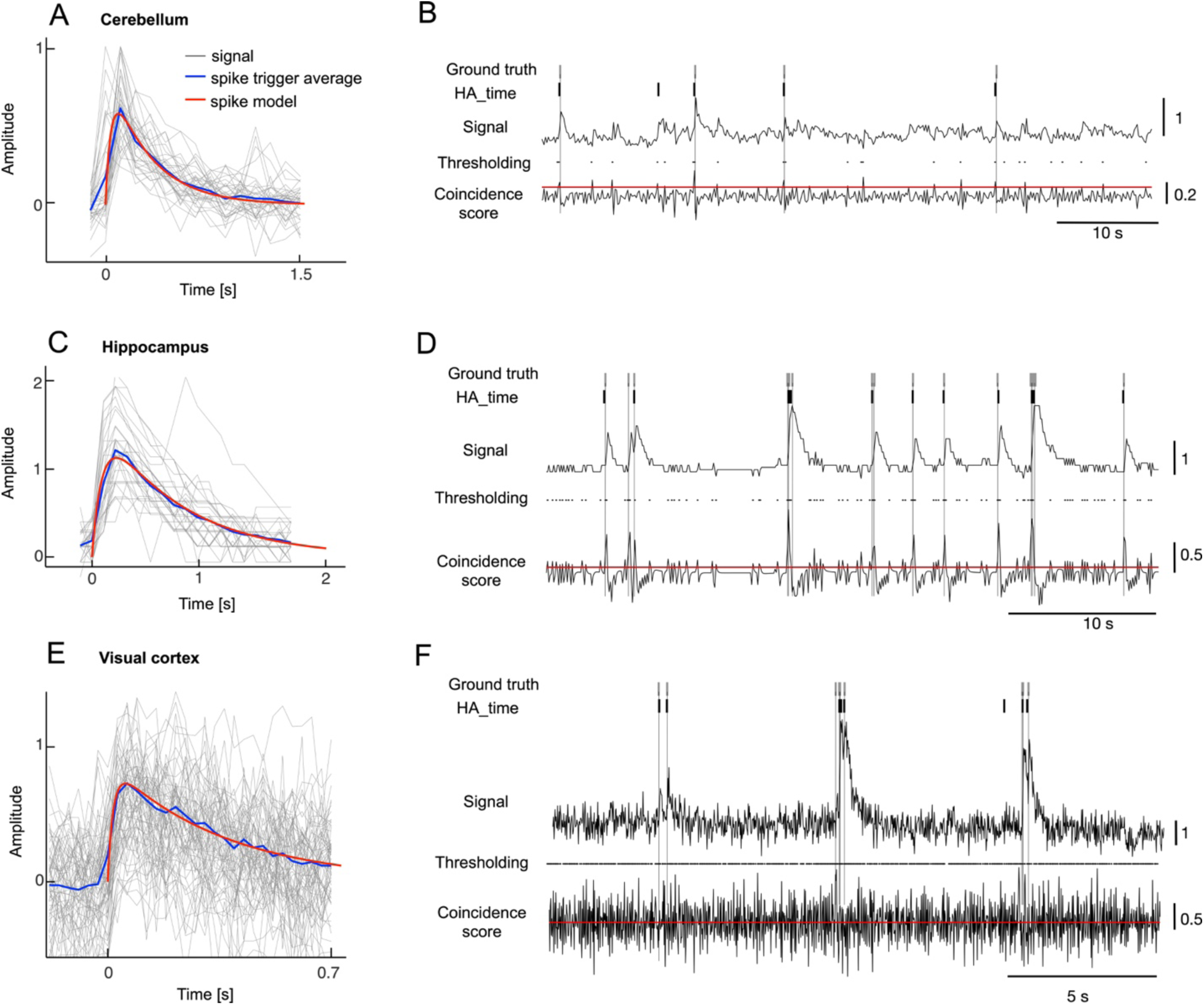
Estimation of the Ca spike model and spike detection by HA_time. A: Ca spike model (red trace, τ_1_ and τ_2_, 0.05 and 0.4 s) and spike-trigger averaged Ca responses synchronized with the onsets of electrical spikes. The Ca spike model and the spike-triggered average were estimated for the thirty-six electrical spikes of five Purkinje cells. Ordinate, amplitude of Ca responses normalized for the peak of the maximum Ca imaging data for the individual cells. Abscissa, time after the onset of the electrical spikes. B: Spike detection by HA_time. The top and bottom traces represent Ca imaging data and the coincidence score of the first-order differential of the Ca imaging data with that of the Ca response model. The candidate spikes detected by conventional thresholding, those estimated by HA_time and the electrical spikes (ground truth) are denoted by black dots, thick and thin bars, respectively. Red lines indicate the thresholds for conventional thresholding. C, D and E, F: similar to A and B but for the hippocampus (τ_1_ and τ_2_, 0.1 and 0.75 s) and visual cortex data (0.01 and 0.2 s), respectively.

Figure 3B illustrates the performance of HA_time in detecting spikes from the Ca imaging data of cerebellar cortex cells. The coincidence thresholding (black dots in Fig. 3B) detected the true spikes (ground truth, gray bars) as well as many false-positive spikes. HA_time effectively selected the true spikes, rejecting many false alarm spikes from the spike candidates. Comparison of the spikes detected by HA_time (dark bars) with the ground truths (gray bars) indicated that HA_time almost perfectly detected spikes and correctly estimated the spike time from the Ca imaging data for the cerebellum (Fig. 3B). HA_time also performed almost completely correctly, rejecting many false alarms detected by conventional thresholding, and it estimated the spike times for the hippocampus and visual cortex data (Figs. 3D and F).

### Nonlinearity analysis of the experimental data sets

We found strong nonlinearity in the Ca imaging data of the hippocampal and visual cortex data during burst activities. Therefore, nonlinearity analysis was conducted by plotting the amplitudes of the Ca imaging data as a function of the linear prediction of the Ca response model for the entire cerebellum, hippocampus and visual cortex data.

The Ca imaging data of the cerebellum roughly agreed with the linear prediction for spike trains (dark and red traces in Fig. 4A), and correspondingly, the regression analysis revealed a fine match between the two (blue line in Fig. 4B, *y = 1.1 x*). Conversely, the nonlinearity analysis of the Ca imaging data revealed significant sub- and superlinearity in the hippocampus and visual cortex, respectively (dark and red traces in Figs. 4C and E). The nonlinearity models were constructed by fitting the plots with logarithmic functions (blue lines in Figs. 4D and F, *y = −3.8 e*^*(−0.16x)*^ *+ 3.9 and y = 0.67 e*^*0.88x*^). The nonlinearity in the hippocampus and visual cortex data was compensated by multiplying the Ca imaging data by the inverse of the nonlinearity models (blue traces in Figs. 4C and E). The compensated Ca imaging data were then fed into HA_time.

**Figure 4:**
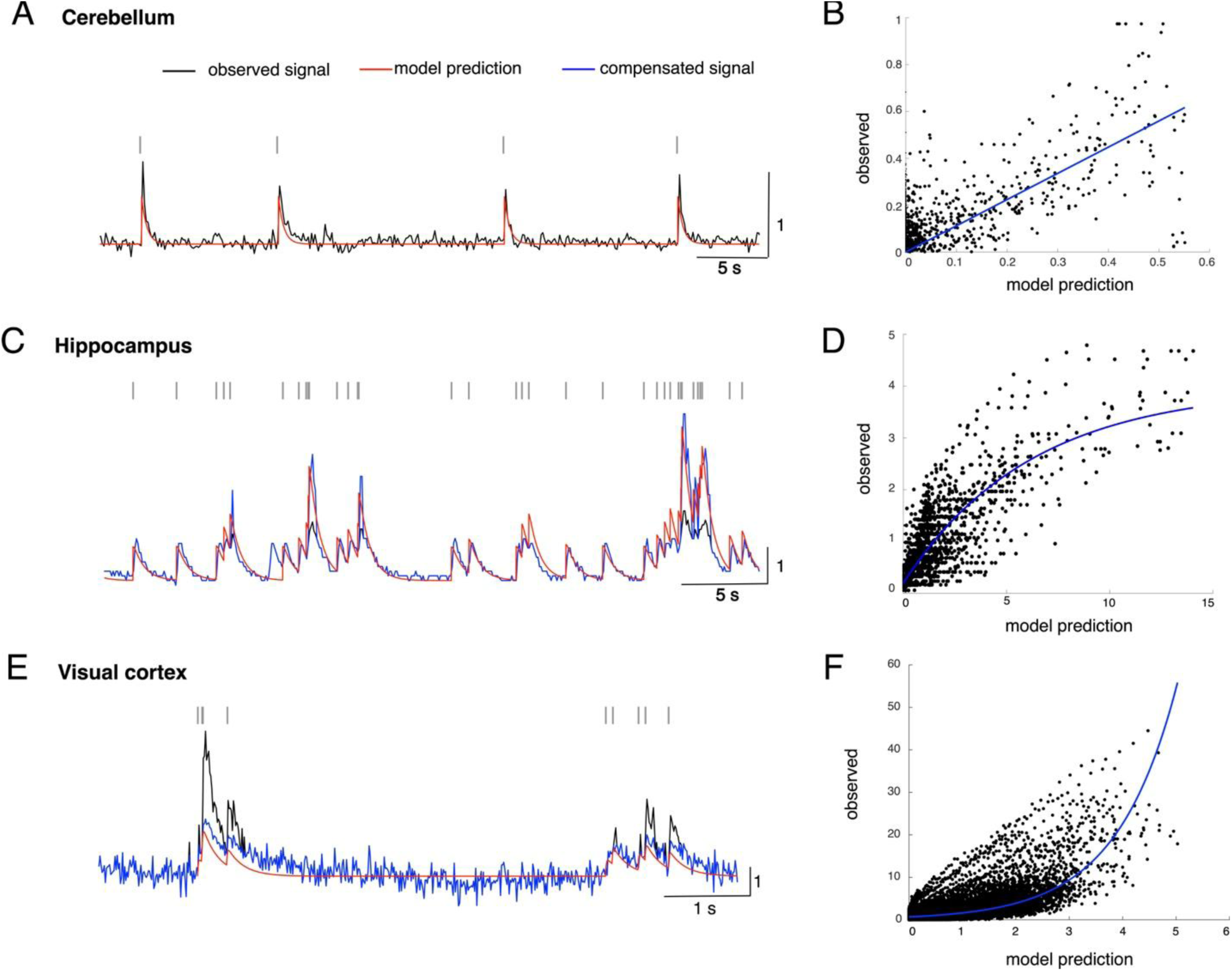
Nonlinearity analysis of Ca imaging data. A, C and E depict the Ca imaging data of the cerebellum, hippocampus and visual cortex data, respectively. Black, red and blue traces represent the observed Ca imaging data, linear prediction of the Ca response model for spike trains and compensated Ca imaging data, respectively. B, D and F depict scatter diagrams for the Ca imaging data in the three experimental data sets as a function of the linear prediction of the Ca response model for spike trains.

### Performance evaluation for experimental data

The performance of HA_time in detecting spikes and in estimating the spike time was studied for the cerebellum, hippocampus and visual cortex data by leave-one-out cross-validation and compared with those of four benchmark algorithms^16,17,22,23^ (cf. Methods).

The spikes detected by HA_time (dark bars) matched fairly well with the ground truths (gray bars) for all of the three experimental data sets. MLspike (red bars) performed fine on the hippocampus data but rather poor on the cerebellum and visual cortex data with many false positives. The remaining three algorithms (yellow, green and purple bars) performed rather poorly for all of the three experimental data sets with many false-positive or missing spikes (Figs. 5A-C).

**Figure 5:**
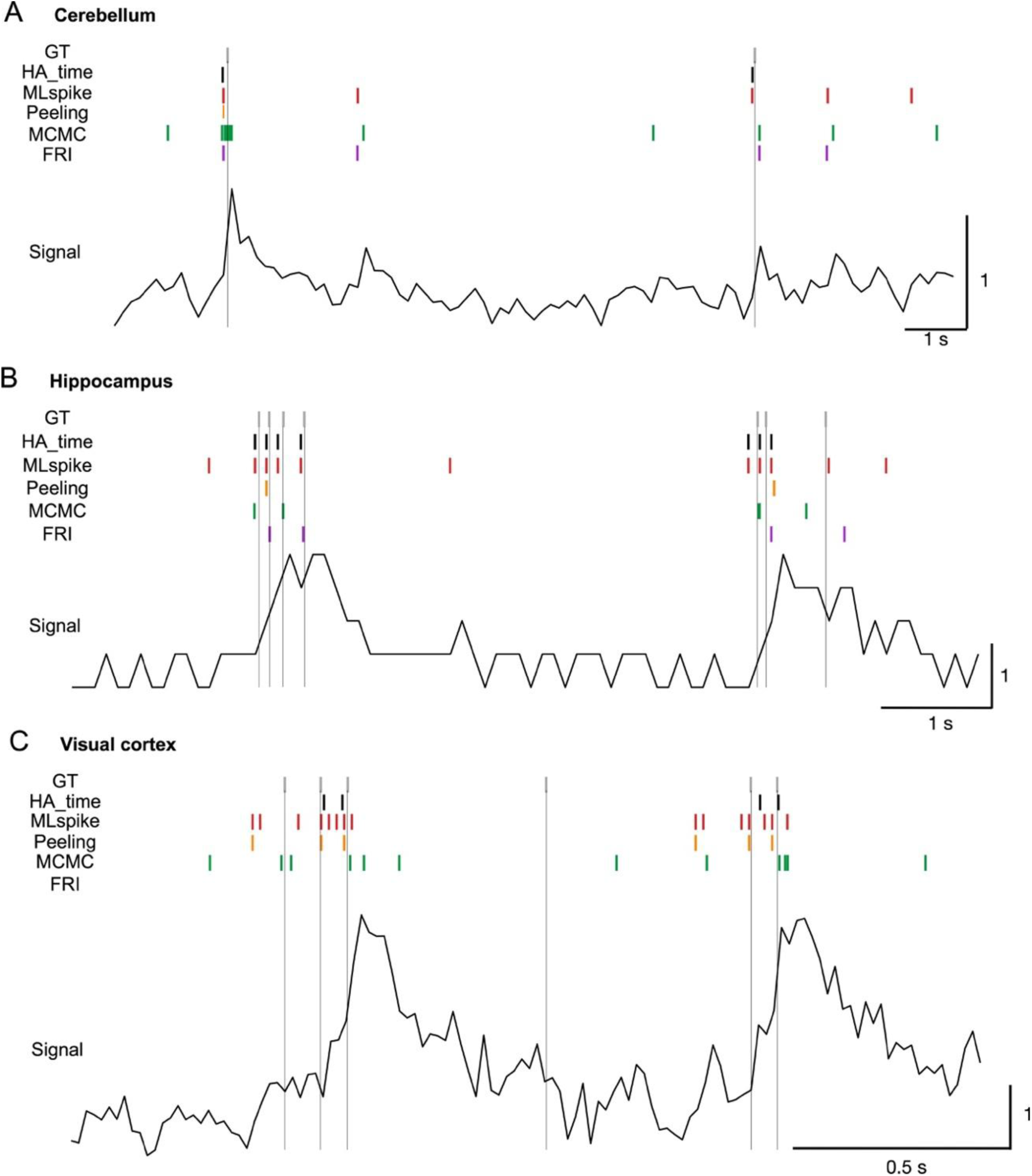
Spike detection by HA_time and benchmark algorithms. A, B and C depict examples of spike detection by HA_time (black bars), MLspike^23^ (red), Peeling^16^ (orange), MCMC^22^ (green) and FRI^17^ (purple) algorithms for the cerebellum, hippocampus and visual cortex data, respectively. Dark traces represent the Ca responses. Thin vertical lines indicate the timing of the ground truth (GT) given by electrical spikes.

We estimated spike detection performance by the F1-score of receiver operating characteristic (ROC) analysis (cf. Methods). Among all of the algorithms, HA_time performed best, with statistical significance in the F1-score for the visual cortex data (0.6 ± 0.04 for HA_time; 0.47 ± 0.05 for MLspike, p = 0.01 for HA_time vs. MLspike; 0.49 ± 0.06 for Peeling, p = 0.01 for HA_time vs. Peeling; 0.39 ± 0.06 for MCMC, p = 0.002 for HA_time vs. MCMC; 0.09 ± 0.07 for FRI, p = 0.002 for HA_time vs. FRI). For the hippocampus data, the superiority of HA_time (0.56 ± 0.11) was also clear, with statistically significant F1-scores compared to those for the benchmark algorithms (0.35 ± 0.08 for Peeling, p = 0.004 for HA_time vs. Peeling; 0.39 ± 0.04 for MCMC, p = 0.02 for HA_time vs. MCMC; 0.14 ± 0.04 for FRI, p = 0.006 for HA_time vs. FRI) except for MLspike (0.53 ± 0.04). However, the significant superiority of HA_time (0.77 ± 0.21) over the benchmark algorithms was limited to Peeling (0.51 ± 0.16, p = 0.02 for HA_time vs. Peeling) and MCMC (0.34 ± 0.13, p = 0.02 for HA_time vs. MCMC) for the cerebellum data. There was no statistical significance in the difference between HA_time and MLspike (0.66 ± 0.31) or FRI (0.53 ± 0.25), probably due to the smaller number of cells (n=5) in the cerebellum data (Fig. 6A).

**Figure 6:**
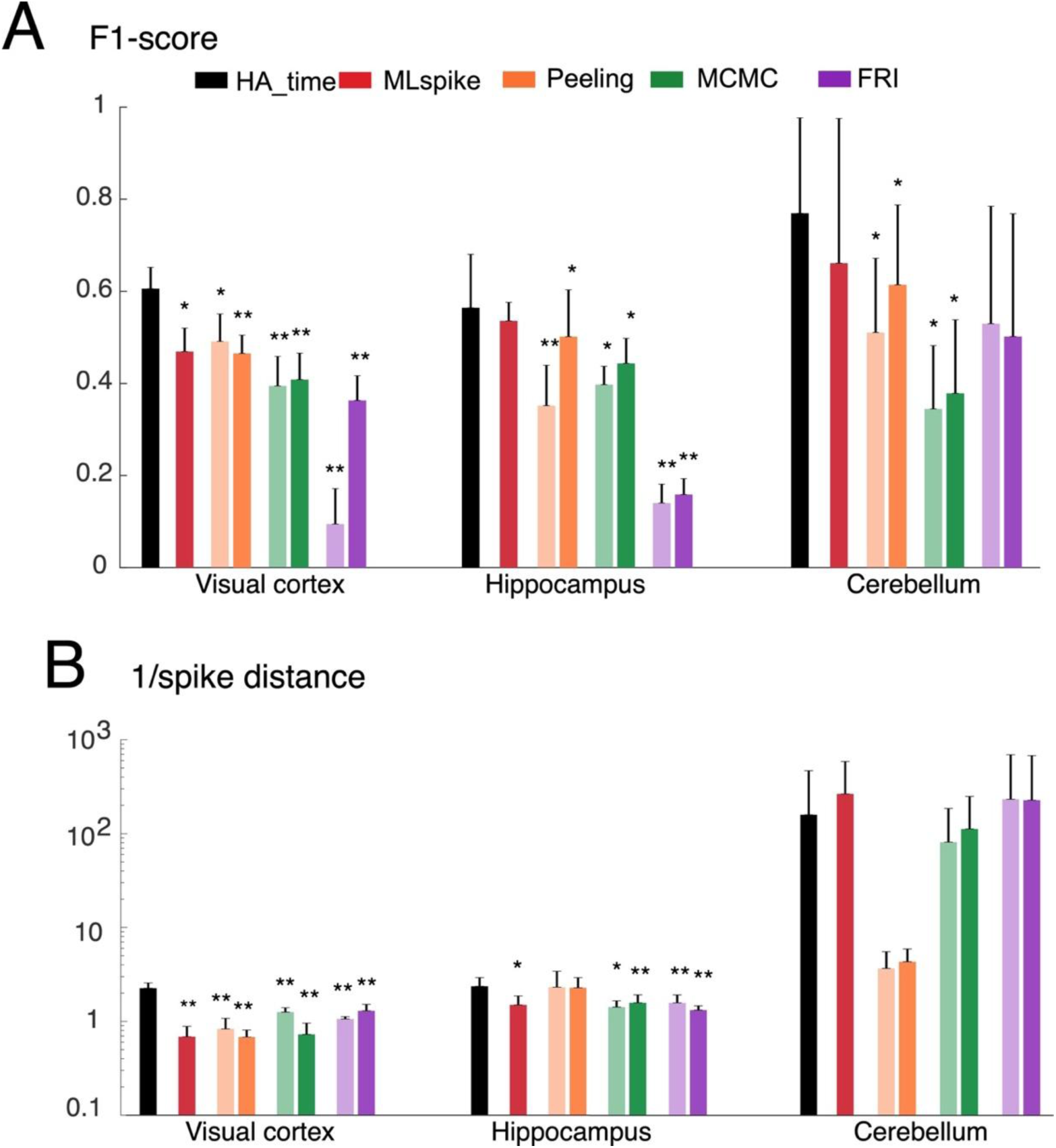
Performance benchmark for experimental data. A and B, F1-score and inverse of spike distance for HA_time (black columns), MLspike (red), Peeling algorithm (orange), MCMC (green) and FRI (purple). The scores of the Peeling, MCMC and FRI algorithms for the original settings and those supplemented with the information provided by the HA_time are shown by dense and faint colors, respectively. The ordinates in A and B are shown in linear and log scale, respectively. The columns represent the mean with error bars of +2SEM. Asterisks indicate significance level by Wilcoxon signed-rank tests between HA_time and the benchmark algorithms. * and ** denote p<0.05 and p<0.01, respectively.

The superiority of HA_time in terms of the precision of spike time estimation was also found by the inverse of the spike distance^31^ (cf. Methods). Among all the algorithms, HA_time performed best with statistical significance over all the benchmark algorithms for the visual cortex data (1/spike distance, 2.2 ± 0.3 for HA_time; 0.7 ± 0.2 for MLspike, p = 0.002 for HA_time vs. MLspike; 0.8 ± 0.2 for Peeling, p = 0.002 for HA_time vs. Peeling; 1.2 ± 0.1 for MCMC, p = 0.002 for HA_time vs. MCMC; 1 ± 0.1 for FRI, p = 0.002 for HA_time vs. FRI). For the hippocampus data, except for Peeling (2.3 ± 1.1), HA_time (2.3 ± 0.5) outperformed the benchmark algorithms with statistical significance (1.5 ± 0.3 for MLspike, p = 0.02 for HA_time vs. MLspike; 1.4 ± 0.2 for MCMC, p = 0.01 for HA_time vs. MCMC; 1.5 ± 0.3 for FRI, p = 0.02 for HA_time vs. FRI). However, for the cerebellum data, no statistical significance was found for HA_time compared with the benchmark algorithms (158.9 ± 309.9 for HA_time; 265.8 ± 325.1 for MLspike; 3.6 ± 1.8 for Peeling; 81.4 ± 104.7 for MCMC; 233 ± 462.1 for FRI, cf. Fig. 6B).

Notably, Peeling, MCMC and FRI were outperformed by HA_time and MLspike, probably because they did not provide an effective routine to precisely estimate the algorithm parameters in the presence of the nonlinearity of Ca responses, which is clearly observed in the hippocampus and visual cortex data. In support of this view, their performance was improved when we provided the parameters of the Ca spike model estimated by HA_time and compensated for the nonlinearity of the Ca imaging data (cf. light and dark bars, Fig. 6A-B).

### Performance evaluation for simulation data

We further investigated the performance of HA_time and the benchmark algorithms by simulating the Ca responses sampled for a broader range of conditions than the experimental ones, including the mean firing frequency of the spike train, the nonlinearity of the Ca responses, the sampling rate of the two-photon recording, the dye dynamics of the Ca responses (time decay constant for the Ca responses) and the SNR (cf. Methods).

The systematic analysis of performance as a function of the experimental parameters revealed that the three parameters, the mean firing frequency, the nonlinearity and the sampling rate, strongly influenced the relative performances of the examined algorithms. In contrast, the remaining two parameters, the dye dynamics and the SNR, altered the performance only in a quantitative manner without significant change in the configuration of the performance changes (Sup Figs. 1 and 2).

Figure 7 illustrates the performance changes as a 3D display by pseudocolor representation as a function of the mean firing frequency and the nonlinearity for three different sampling rates (τ_2_ and SNR were fixed at 0.2 s and 5, respectively). HA_time outperformed all of the benchmark algorithms in terms of the F1-score across the entire parameter range (Fig. 7A), except for statistically nondiscriminable MLspike under the high sampling rate and low firing frequency condition (30-60 Hz and <=2 Hz). The three other algorithms, Peeling, MCMC and FRI, exhibited feasible performance only under the high sampling rate, low firing frequency and weak nonlinearity (α = 0.6-1) condition.

**Figure 7:**
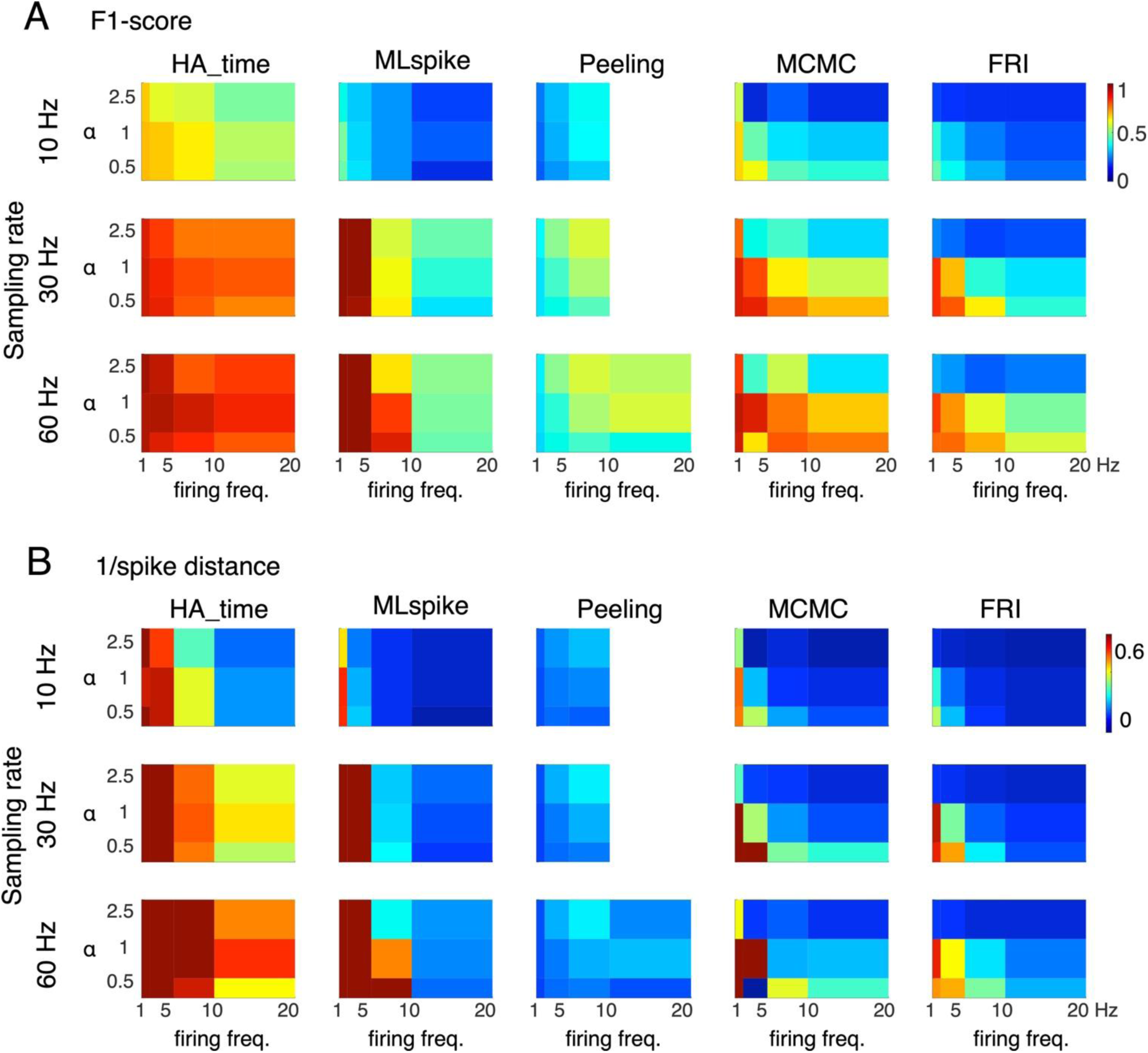
Performance of HA_time and benchmark algorithms for simulation data. A and B, Pseudocolor 3D maps of F1-score and inverse of spike distance as a function of mean firing frequency (abscissa) and nonlinearity (α, ordinate) for the three different sampling rates (10, 30 and 60 Hz). τ_2_ and SNR were fixed at 0.2 s and 5, respectively. Blank areas indicated that the Peeling algorithm failed to perform in a sufficient time for the cases of high firing frequency and low sampling rate.

The performance in the temporal precision of spike timings, estimated as the inverse of the spike distance, also showed the same tendency as that for the F1-score. MLspike performed best under the high sampling rate and low firing frequency condition, and the other three benchmark algorithms exhibited feasible performance only under the low firing frequency, weak nonlinearity and high sampling rate condition. Conversely, HA_time outperformed all benchmark algorithms over the entire parameter range except for under the high sampling rate and low firing frequency condition (Fig. 7B).

Figure 8 shows profiles of the F1-scores and the inverse of the spike distance values for low (10 Hz) and high sampling rates (60 Hz) as a function of the mean firing frequency (the nonlinearity parameter α was fixed at 1). HA_time outperformed all benchmark algorithms under the low sampling rate condition across the entire range of firing frequencies (1-20 Hz, Figs. 8A and B). The superiority of our algorithm over the benchmark algorithms was also found under the high sampling rate condition across the entire range of spike frequency, except for MLspike, which performed best under the low firing frequency (<= 2 Hz) condition. However, the performance of MLspike reduced to the level of the other benchmark algorithms as the firing frequency increased (Figs. 8C and D).

**Figure 8:**
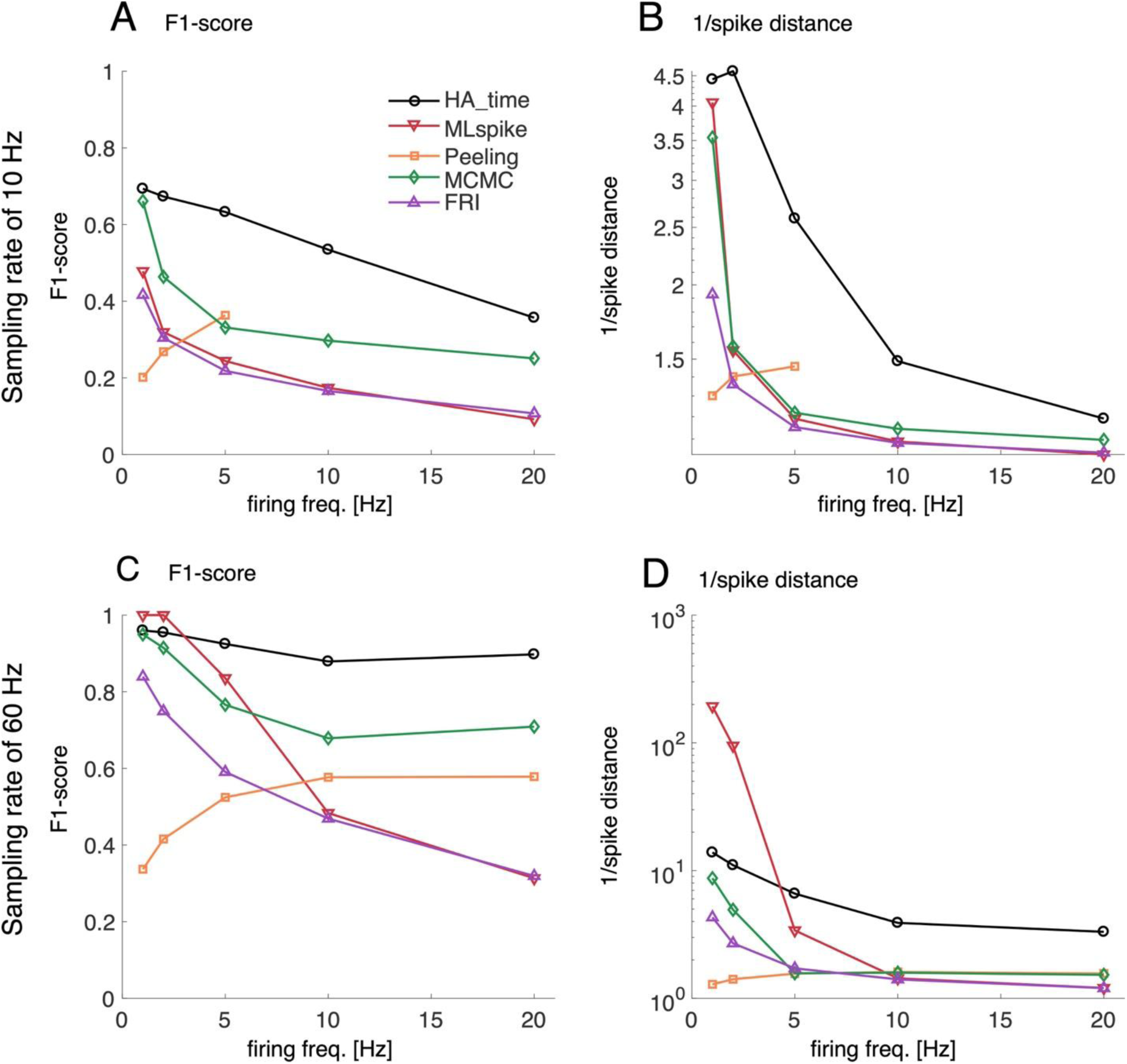
Firing frequency profiles of performance. A and B, F1-scores and inverse of spike distance values for HA_time and the benchmark algorithms as a function of firing frequency at a sampling rate of 10 Hz. C and D, those at 60 Hz. The nonlinearity parameter α was fixed at 1. The color conventions for the algorithms are the same as those in Fig. 6.

## Discussion

HA_time aimed to resolve two challenging issues, reliable spike detection and high spike time precision in the presence of the nonlinearity, slow dynamics and low SNR of Ca imaging data. The difficulty in achieving this goal arose from the spike dynamics of the target neurons and/or kinematics of the Ca responses that may vary dramatically across brain regions. The slow Ca kinematics and the low SNR may limit the temporal precision of the information conveyed by the Ca imaging data. HA_time overcame this difficulty by combining a generative model for the nonlinearity and Ca dye dynamics with a supervised classifier. It estimated the Ca spike model by Bayesian inference assuming the size and shape constancy of the spike, compensated for the nonlinearity of the Ca responses by nonlinearity analysis, and detected the spikes from the compensated Ca imaging data using the ground truths as supervised information. Hyperacuity precision of spike time estimation was achieved by recalibrating the spike time to minimize the residual errors in the Ca response model prediction using the hyperacuity vernier scale. The combined approach may improve the performance of HA_time in two ways. The Ca response model helped spike detection as well as spike time estimation with higher temporal precision than the sampling resolution, while the supervised learning compensated for fluctuations in the Ca imaging data due to the noise and sampling jitters that are not considered by the generative model.

We also developed the generative-model algorithm, hyperacuity Bayes, (see Methods and Supplemental Information) which is a partial algorithm of HA_time. In this algorithm, estimation of the generative model by Bayesian inference with ground-truth information was very robust. However, we ascertained the superiority of the combination approach (i.e. HA_time) in estimating spike timing over the one maximizing the likelihood estimate of the generative algorithm (i.e. hyperacuity Bayes) by the significantly higher F1-score as well as the inverse of the spike distance for the hippocampus and visual cortex data (Fig. S3). Regarding temporal precision, compared with conventional thresholding, HA_time reliably improved up to 20-fold for cases where the sampling rate of Ca imaging was as high as 40-60 Hz. Rapid progress in the development of two-photon imaging techniques with higher sampling rates and faster Ca kinematics may reinforce the superiority of HA_time over conventional methods. We also found that HA_time outperformed the other four hyperacuity algorithms in terms of the F1-score, with feasible statistical significance in the visual cortex and hippocampus data that include a relatively large number of cells. However, for the cerebellum data, in which the number of cells is rather small, HA_time outperformed only the Peeling and MCMC algorithms. HA_time also outperformed all benchmark algorithms in terms of the precision of spike time estimation, which was estimated as the inverse of the spike distance, with statistical significance in visual cortex data. However, HA_time outperformed only MCMC and FRI in the hippocampus data, and outperformed none of the algorithms in the cerebellum data.

We conducted a systematic study of algorithm performance on simulation data that covered a broader range of parameters than the experimental conditions, including the mean neuronal firing frequency, the nonlinearity and relaxation time of the Ca dyes and the sampling rate. The performances of the F1-score as well as the inverse of the spike distance values studied as the functions of those parameters pointed to the mean firing frequency, the nonlinearity and the sampling rate as the most important parameters that influenced the changes in performance. In contrast, the SNR and the relaxation time of the dyes are the less important parameters that only influenced the size while not significantly changing the shape of the performance functions. The F1-score and the inverse of spike distance functions highlighted the superiority of HA_time over the other algorithms, showing high scores of spike detection and high spike time precision across the entire range of parameters except for the high sampling rate and low firing rate condition, where MLspike slightly outperformed HA_time. The performance of the other benchmark algorithms remained feasible only under the weak nonlinearity, high sampling rate and low firing frequency condition. Conversely, HA_time maintained high performance under the strong nonlinearity and/or high firing frequency conditions frequently encountered in the cortical cells of behaving animals.

The simulation analysis of the performance for HA_time and the benchmark algorithms across a wide range of experimental conditions for two-photon recordings may provide useful information for selection of the best algorithm for given experimental conditions. Although the application of HA_time is limited to cases where ground-truth signals are available, it may also be applicable to cases where simultaneous electrical recordings are unavailable as follows. One may estimate the parameters presently studied for the experimental data of a two-photon recording using a maximum likelihood method, then train HA_time with a newly generated simulation data including spikes for the estimated parameters, and finally use the trained HA_time for spike estimation in the experimental data of the two-photon recording. This approach may benefit from the combination of the generative and supervised approaches, as shown by the present study. In summary, HA_time is useful to improve spike detection and temporal precision in spike time estimation across a wide range of the experimental conditions for two-photon recording in cases where examples of simultaneous ground-truth signals (electrical spike recording) are available.

## Methods

### Hyperacuity support vector machine (HA_time)

HA_time detects spikes contained in the Ca responses of two-photon recordings in three steps: 1) estimation of the Ca response model by the expectation-maximization (EM) algorithm assuming the shape and size constancy of the spike model, 2) spike detection in the Ca responses by a support vector machine (SVM) assisted by the information of the Ca response matching with the Ca spike model, and 3) spike time estimation for the detected spikes to minimize the errors between the Ca response model prediction and the Ca imaging data using the vernier scale ten-fold finer than the sampling interval of two-photon recordings.

#### Ca spike model estimation by Bayes inference

We estimated the parameters of the Ca response model assuming that all of the Ca responses in the two-photon recording originate from a unique Ca spike model *g(t, T*, τ) and vary due to the noise and sampling jitters (SJs)^19^.

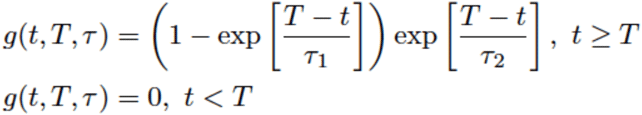

where t, T, τ = (τ_1_, τ_2_) are the time, the spike onset, and the rise and decay time constants, respectively.

We estimated a Ca response model whose parameters are the model amplitude (a), baseline (b_0_) and noise (σ) using the EM algorithm, while the time constants of spike response (τ) were estimated by iterative alternate coordinate 1D grid search (see Supplemental Information).

#### Spike detection by SVM

We conducted spike detection and spike time estimation by an SVM supplemented with the information from the Ca response model to improve the performance of the SVM. We estimated the coincidence scores, determined as the convolution (*dy/dt * dg/dt*) of the first-order derivative (*dy/dt*) of the Ca signals of a 2-photon recording with that (*dg/dt*) of the Ca response model (*g)* estimated by Bayesian inference for the training data, and sampled data segments that exceeded the threshold as the spike candidates. The threshold and the length of data segments (the number of data points before and after the point exceeding the threshold) were optimized according to the F1-score for the training data (cf. cross-validation in Methods). The SVM was trained to classify the spike candidates into spikes or non-spikes by feeding the spike candidates and the coincidence scores as the primary and attribute inputs, respectively, and the electrical spikes (ground truth) as the teaching signals.

#### Hyperacuity spike time estimation

The trained SVM was used for spike detection in the test data. The pseudo-spike times (PTs) were tentatively determined for the detected spikes as those for which the coincidence score exceeded the threshold. We assumed that the PTs may vary due to the SJ (difference between pseudo- and true spike time) and estimated the SJ to minimize the prediction errors between the Ca response and the Ca response model by systematically changing the SJ according to a vernier scale 10-fold finer than the sampling interval. The true spike time (TT) was calculated as the sum of PT and SJ (cf. the inset of Fig. 1C). We subtracted the trace of the preceding spike from the succeeding one for spike time estimation in cases where the preceding spikes overlap with the succeeding ones.

### Other benchmark algorithms

We evaluated the performance of four hyperacuity algorithms: the MLspike algorithm^23^, the Peeling algorithm^16^, the finite-rate innovation method^17^ (FRI), and the Monte Carlo Markov chain method^22^ (MCMC). HA_time and MLspike used the ground-truth signals, given as the electrical spikes for algorithm optimization, whereas the remaining three algorithms did not. We studied how the performance of the three algorithms may be improved in cases where they are supplemented with our parameter settings for the Ca response model and nonlinearity (cf. ^16,17,22^). To compute the temporal improvement in the hyperacuity algorithms, we also conducted a conventional thresholding algorithm, whose threshold was optimized in the range of 0-4 SD by maximizing the F1-score of the training data.

### Experimental data sets

We collected simultaneous recordings of electrical and two-photon recording of the Ca signals in three cortical areas (cerebellar, hippocampal and visual cortices) using three different calcium dyes as described below.

#### Recording of cerebellar Purkinje cell complex spikes

We collected experimental data for the complex spikes of five cerebellar Purkinje cells from the work of ^32^, where simultaneous two-photon Ca imaging (sampling rate, 7.8 Hz) using multicell bolus loading of Cal-520 dye and extracellular recording (sampling rate, 20 KHz) was performed on adult mice.

#### Recording of hippocampal CA3 neurons

We collected the simultaneous cell-attached recording (sampling rate, 20 KHz) and one-photon images (10 Hz) of Ca responses from nine CA3 pyramidal neurons in organotypic cultured slices of rats stained with OGB-1AM dye^33^. The Ca signals were normalized by the peak of the Ca spike model estimated for individual cells.

#### Recording of the primary visual cortex

We collected the data from the paper of ^30^. The data set contained simultaneous loose-seal cell-attached patch recordings (sampling rate, 20 KHz) and 2-photon recordings of Ca responses from eleven GCaMP6f-expressing neurons in a behaving mouse visual cortex (sampling rate, 60 Hz).

### Simulation data

We conducted a simulation of the Ca responses for the three experimental data sets: those of the cerebellar, hippocampal and primary visual cortices. Spike events were generated according to a Poisson distribution whose mean firing rate varied across 1-20 Hz. The Ca responses were simulated by convolving the double exponentials with time constants for rise and decay with the spike events. The rise time constant τ_1_ was fixed at 0.01 s, while the decay time constant τ_2_ was varied across 0.2-1 s, corresponding to those for the OGB-1, Cal-520 and GCAMP6f dyes. We introduced the parameter α to reproduce the nonlinearity found in the Ca responses f(t) in the three cortices as

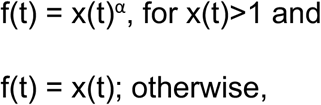

where x(t)=g(t)*s(t) is the linear response of the Ca spike model given the spike train s(t) and the parameter α for saturation (α<1) and superlinearity (α>1) was varied in the range of 0.2-3, corresponding to the values found in the three experimental data sets. Finally, Gaussian noise was added to reproduce the SNR (3, 5, 10) of the experimental data. For each set of simulation parameters, 500 spike signals in a total of ten cells were generated, and those of five cells were used as the training and test data sets.

### Performance analysis

For evaluation spike detection performance, the correct hit case was defined as that where the time difference of the estimated spike from the true one was smaller than a window of half the sampling interval, and vice versa for the missing case, and the false-positive case was defined as that where the time difference of the true spike from the estimated one was greater than the time window. For data sets with a high sampling rate (30-60 Hz), the window was relaxed to 50 ms.

Receiver operating characteristic (ROC) analysis was conducted for these cases as:

- Sensitivity = Hit / (Hit + misses)
- Precision = Hit / (Hit + False positive)
- F1-score = 2 × (Sensitivity × Precision) / (Sensitivity + Precision)

We estimated the temporal precision of spike time estimation as the inverse of the spike distance, defined as the minimal cost for reconstructing the true spike train from the estimated one, allowing 1 each for deletion or insertion of the spike event and the weighted cost for the shift in the spike time^31^. The spike distance was further normalized by the number of ground-truth spikes. For evaluation of temporal precision improvement, spike time errors were estimated as the absolute time difference between the closest true and reconstructed spikes for all spikes in both spike trains. The hyperacuity improvement was determined as the ratio of the mean spike time errors of conventional thresholding to that of the hyperacuity algorithms.

### Statistical analysis

Performance analysis of all algorithms on the experimental data was conducted by the one-leave-out cross-validation, where the data of one cell and that of the remaining cells were used for testing and training, respectively.

All of the performance scores were estimated as the mean ± 2SEM. To assess statistical significance, we compared the performance of HA_time to that of the benchmark algorithms by a one-sided Wilcoxon signed-rank test and reported the significance level p.

### Hyperacuity Bayesian Algorithm

We also developed a hyperacuity Bayesian (HB) algorithm by reproducing a basically similar algorithm to that for HA_time. HB is applicable for cases where no ground-truth signals are available, maximizing the likelihood for the Ca signals recorded by two-photon recordings. For cases where the training data with ground-truth signals is provided, HB optimized the model parameters to improve the overall performance of spike time inference (see Supplemental Information).

## Data availability

The MATLAB® implementation of our algorithm can be found online (https://github.com/hoang-atr/HA_time). The hippocampal and cerebellar cortex data sets used in this work are available from the authors upon reasonable requests.

## Acknowledgements

This research was conducted under contract with the National Institute of Information and Communications Technology, entitled “Development of network dynamics modeling methods for human brain data simulation systems”. HH and KT were partially supported by the ImPACT Program of Council for Science, Technology and Innovation (Cabinet Office, Government of Japan). HH, YI, and KT were partially supported by JST ERATO (JPMJER1801, “Brain-AI hybrid”).

## Supplemental Information

### Hyperacuity Bayesian algorithm

We also developed a hyperacuity Bayesian (HB) algorithm for spike detection and spike time estimation, maximizing the estimate likelihood for the cases where ground-truth signals are not available. The HB algorithm included supervised and unsupervised versions. The supervised version reproduces essentially similar procedures, such as estimation of the Ca response model, spike detection and spike time estimation using the model information, to those for HA_time by the Bayesian algorithm. Probabilistic models and the EM algorithm are described in the first two sections, and detailed procedures of the two versions of the HB algorithm are described in the next two sections.

#### Data structure and probabilistic model

Let us suppose that K data segments were sampled from data by thresholding while leaving the rest of the data (y_rest_).

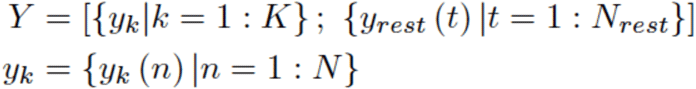

where y_k_(n) is the sampled data at the sampling time t_k,n_= (n-1)dt_0_+t_k,1_ of the k-th window and dt_0_ is the sampling step of the observed data with the sampling frequency f_0_ = 1/dt_0_.

A probabilistic model for spike states is given by

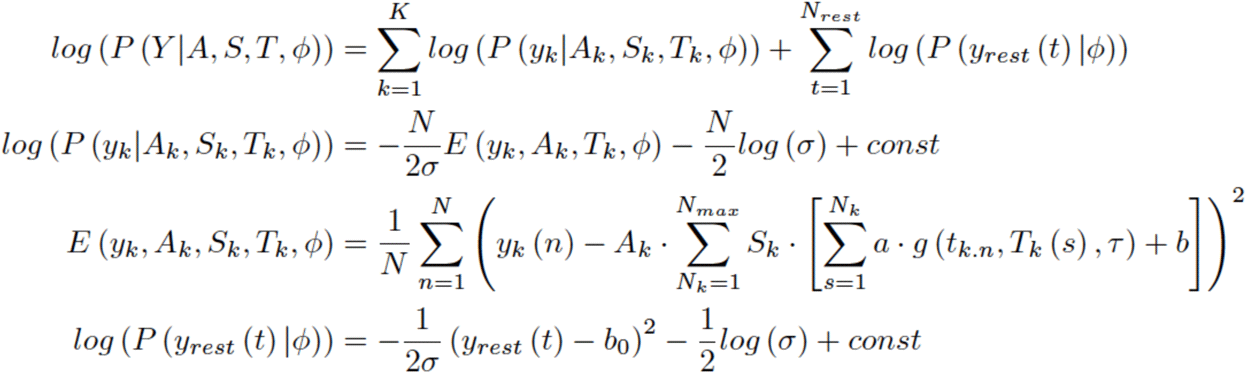

where A_k_ is a spike indicator variable, which represents the presence or absence of spikes in the k-th window, and takes a binary value (0 or 1). S_k_ represents a spike state in the k-th window and takes a binary vector value (Potts spin variable)

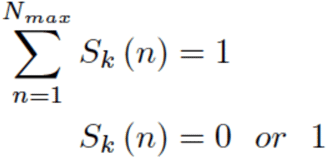

S_k_(N_k_) = 1 means that there are N_k_ spikes in the k-th window. The maximum number of spikes in a window is assumed to be N_max_. T_k_ is a set of spike times in the k-th window, and T_k_(s) is the onset time of the s-th spike. *a* represents the amplitude of the spike response function. *b* and *b*_*0*_ represent the bias in spike and no spike regions, respectively. σ is the variance of the Gaussian noise. A set of global parameters that is assumed to be common for all spikes and the rest of the data is denoted by *ϕ=*(*a, τ,b,b*_0_, *σ*).

We assume the hierarchical noninformative priors as

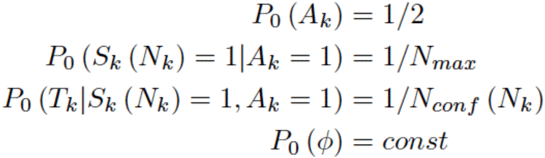

where N_conf_(N_k_) represents the number of configurations of T_k_ for the N_k_ spike case.

#### Expectation maximization (EM) algorithm

In the E-step, the posterior probability of the spike state for the current estimate of the model parameters *ϕ=*(*a, τ,b,b*_0_, *σ*).for combination of the log-likelihood, the noninformative prior and the joint probability for a spike state in the k-th window is given by:

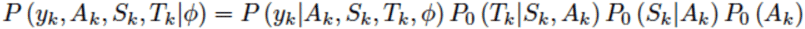

and marginal probability is given by:

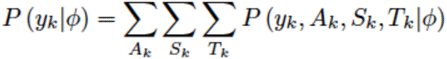

The posterior probability for a spike state is then calculated as

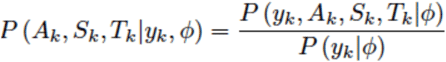

In the M-step, the model parameters were updated to a new value *ϕ*_*new*_ by maximizing the Q-function, defined as

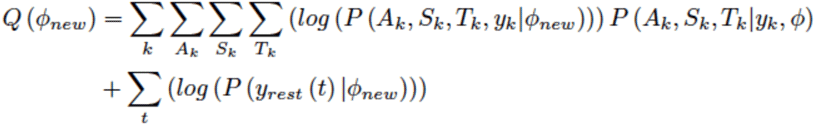

by solving the maximum condition

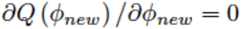

#### Spike estimation for the supervised version of HB

We first extracted continuous regions in which the signals exceed the threshold. Here, the threshold was optimized to maximize the true-positive cases and minimize the false-positive cases referring to the ground truth given by the electrical spikes. Each region is then segmented into fixed-length data segments of 8 points. We divided two overlapping data segments, whose onset intervals were less than the length of the data segment (8 points), into three divided nonoverlapping segments. For example, two segments of points #1-10 and #5-14 were divided into three nonoverlapping segments of points #1-4, #5-10 and #11-14.

Next, we utilized the training data to estimate the model parameters and the classifier for posterior probability. The spike model parameters including the spike amplitude (a), biases (b, b_0_) and noise variance σ were estimated by the EM algorithm (see above), while time constants of the spike response (τ) were estimated by iterative alternate coordinate 1D grid search because the log-likelihood with respect to τ is highly nonlinear. The maximum number of spikes contained in single data segments was estimated for the 95-percentile value of the spike number histogram of the training data segments. The posterior probability of the spike state for training data was estimated for each data segment assuming that they are independent of each other. For overlapping segments, posterior probabilities were integrated among the overlapping segments by Bayes inference. We used a multinomial classifier to predict the number of spikes from the posterior probability.

Finally, we estimated the spike number and spike onset time for test data. The data segments were sampled for the test data by thresholding whose threshold was optimized for the training data, and the posterior probability of the spike state for each segment was calculated in the same way as for the training data. The number of spikes was estimated from the posterior probability for the number of spikes using the multinomial classifier trained for the training data, and spike onset times were estimated with the hyperacuity time step by maximizing the log-likelihood for the estimated number of spikes. The contributions of preceding spikes were subtracted from the data signal, as was done for HA_time.

#### Spike estimation by the unsupervised version of HB

The unsupervised version of HB conducted the spike estimation in essentially the same way as that for the supervised version, except for the algorithm used to estimate the spike state. We conducted Bayesian inference assuming that the initial spike state for each data segment contains only one spike whose waveform represents the spike response. We optimized the threshold and τ = (τ_1_, τ_2_) that maximize the log-likelihood and estimated the number of spikes for each data segment that gave the maximum posterior probability for the given number of spikes.

## Supplemental Figures

**Supplementary Figure 1:**
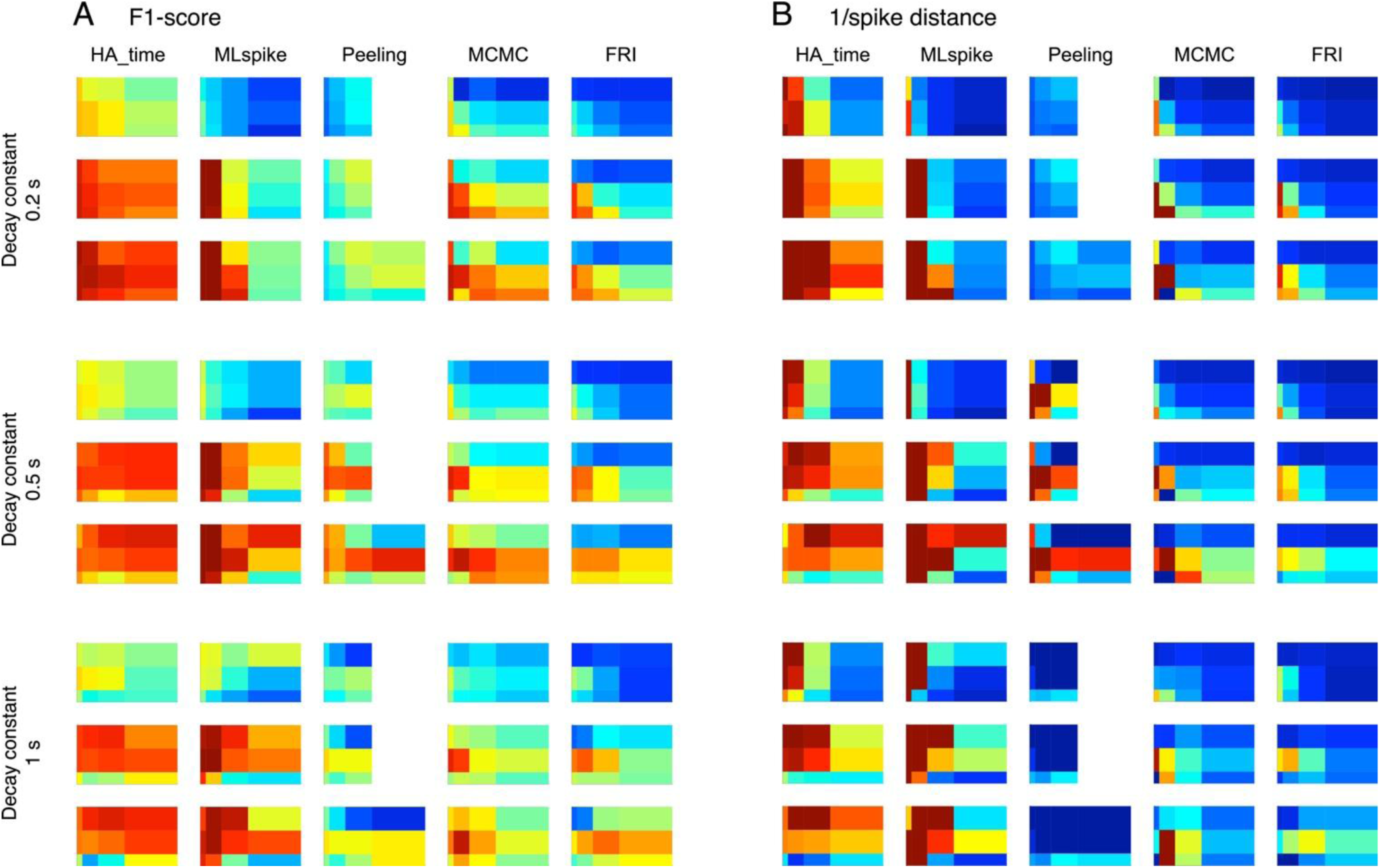
Performance of HA_time and benchmark algorithms on simulation data for variation in the decay time constant of the Ca response model. A and B, performance of F1-score and inverse of spike distance for three different decay time constants compared to that in Fig. 5 for the Ca response model (0.2, 0.5 and 1 s) and three different sampling rates (10, 30 and 60 Hz). The ordinate, abscissa and calibration scale of the pseudocolor maps are the same as those in Fig. 5.

**Supplementary Figure 2:**
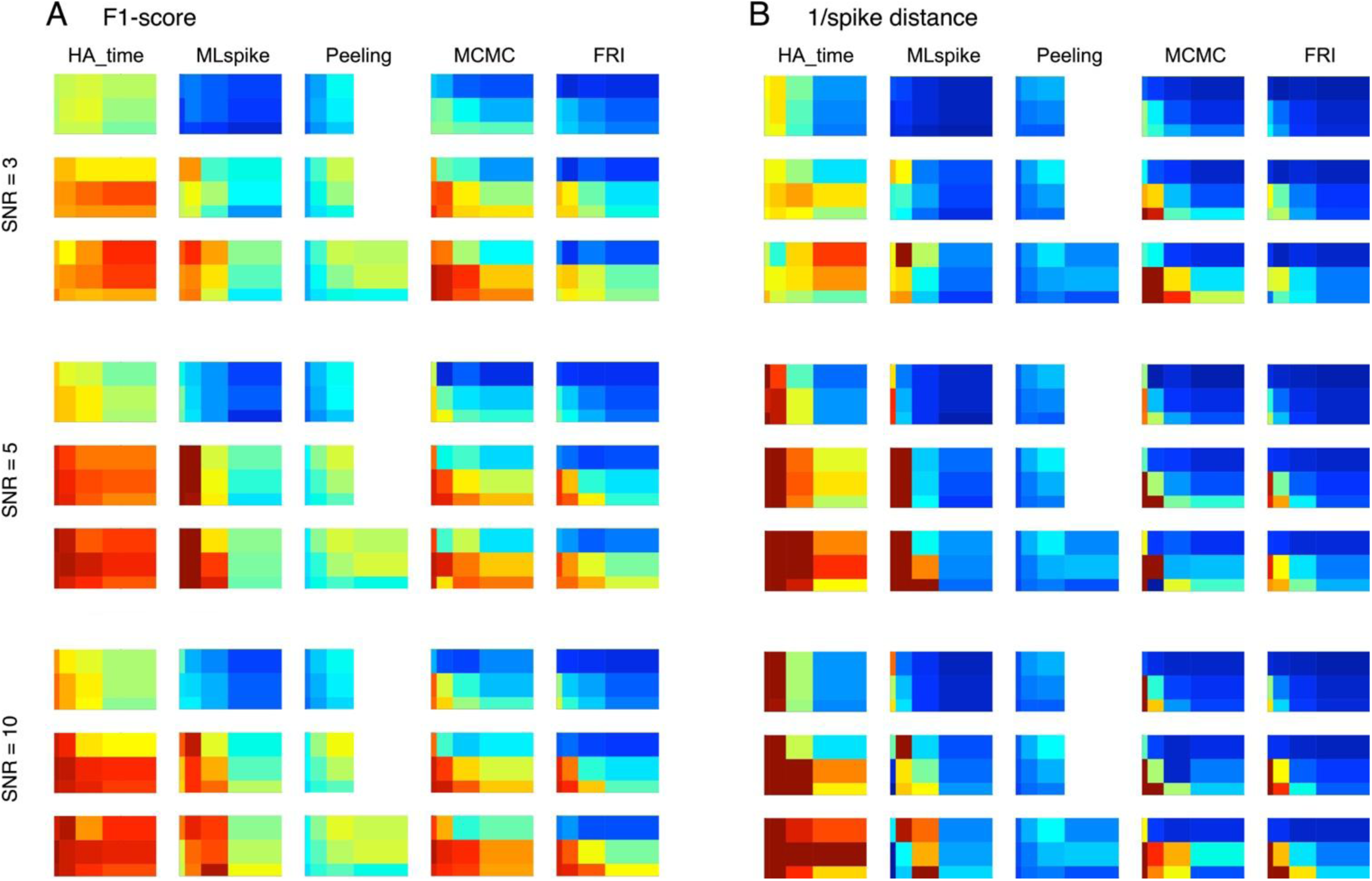
Performance of HA_time and benchmark algorithms on simulation data for variation in the SNR of Ca response signals. A and B, performance of F1-score and inverse of spike distance similar to Fig. S1 but for three different SNRs of the Ca response signals.

**Supplementary Figure 3:**
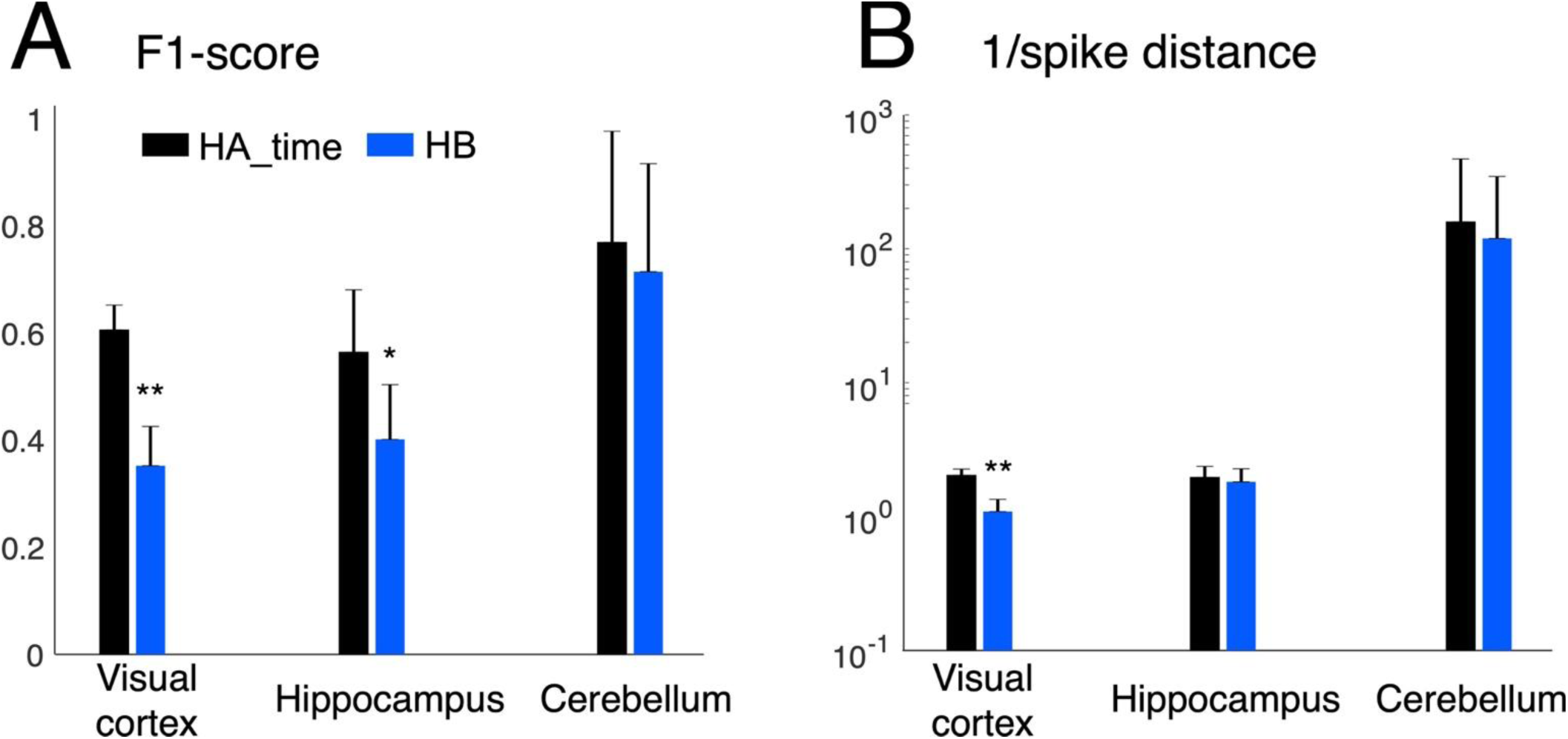
Performance benchmark of HA_time and HB for experimental data. A and B, F1-score and inverse of the spike distance for HA_time (black columns) and HB (blue). Conventions are the same as those in Fig. 6.

